# Extrinsic polarity cues control lamination versus cluster-based organisation in vertebrate retinal development

**DOI:** 10.1101/2025.11.12.688026

**Authors:** Christina Schlagheck, Xenia Podlipensky, Cassian Afting, Ronald Curticean, Irene Wacker, Rasmus R. Schröder, Venera Weinhardt, Lucie Zilova, Joachim Wittbrodt

## Abstract

Photosensitive organs are essential for most animals to perceive and respond to their environment. While the gene regulatory networks establishing retinal identity are deeply conserved across metazoans (reviewed in Gehring, 2012; Vopalensky & Kozmik, 2009; Hahn et al., 2023), the retinal architecture varies widely—from invertebrate compound eyes to vertebrate camera-type eyes (Lamb et al., 2007; Schwab, 2017; Arendt & Wittbrodt, 2001).

Despite this morphological diversity, early eye anlagen in both, invertebrates and vertebrates, share an initial pseudo-stratified epithelial organization (Weasner & Kumar, 2022; Randlett et al., 2010; Das et al., 2003; Kitambi & Malicki, 2008), which is maintained and elaborated into multi-layered retinae in vertebrates. In contrast, the invertebrate neuroepithelium is re-organized as ommatidia develop. Laminar organisation of the vertebrate retina appears to be a consequence of initial polarisation of the retinal neuroepithelium. This is, however, challenging to test in the organismal context.

To address the plasticity of retinal architecture and the impact of epithelial polarity on the structuring of retinal tissue, we take advantage of retinal organoids derived from medaka (*Oryzias latipes*) (Zilova et al., 2021) that allow to modulate polarity cues and test their impact on the level of epithelialization and structural organisation of the forming retina.

We show that under specific culture conditions, medaka retinal organoids undergo a striking morphological switch depending on the level of apico-basal polarity imposed. When polarity cues are continuously provided, a laminated retinal epithelium is established in the organoid. The absence of polarity cues results in the formation of horizontal cellular clusters containing the retinal cell types, which form the vertical retinal column in the developing embryo.

We demonstrate that the emergence of this alternative retinal architecture is associated with a loss of epithelial polarity, notably the absence of extracellular matrix (ECM) components, such as laminin, which efficiently rescues lamination. Our findings indicate that tissue-level polarization and lamination in vertebrate retinae require specific extrinsic cues, and that in their absence, differentiating retinal cell types self-organize into structurally distinct, retinal units. This reveals an unexpected plasticity in vertebrate retinal development and indicates a potential for alternative modes of retinal patterning.

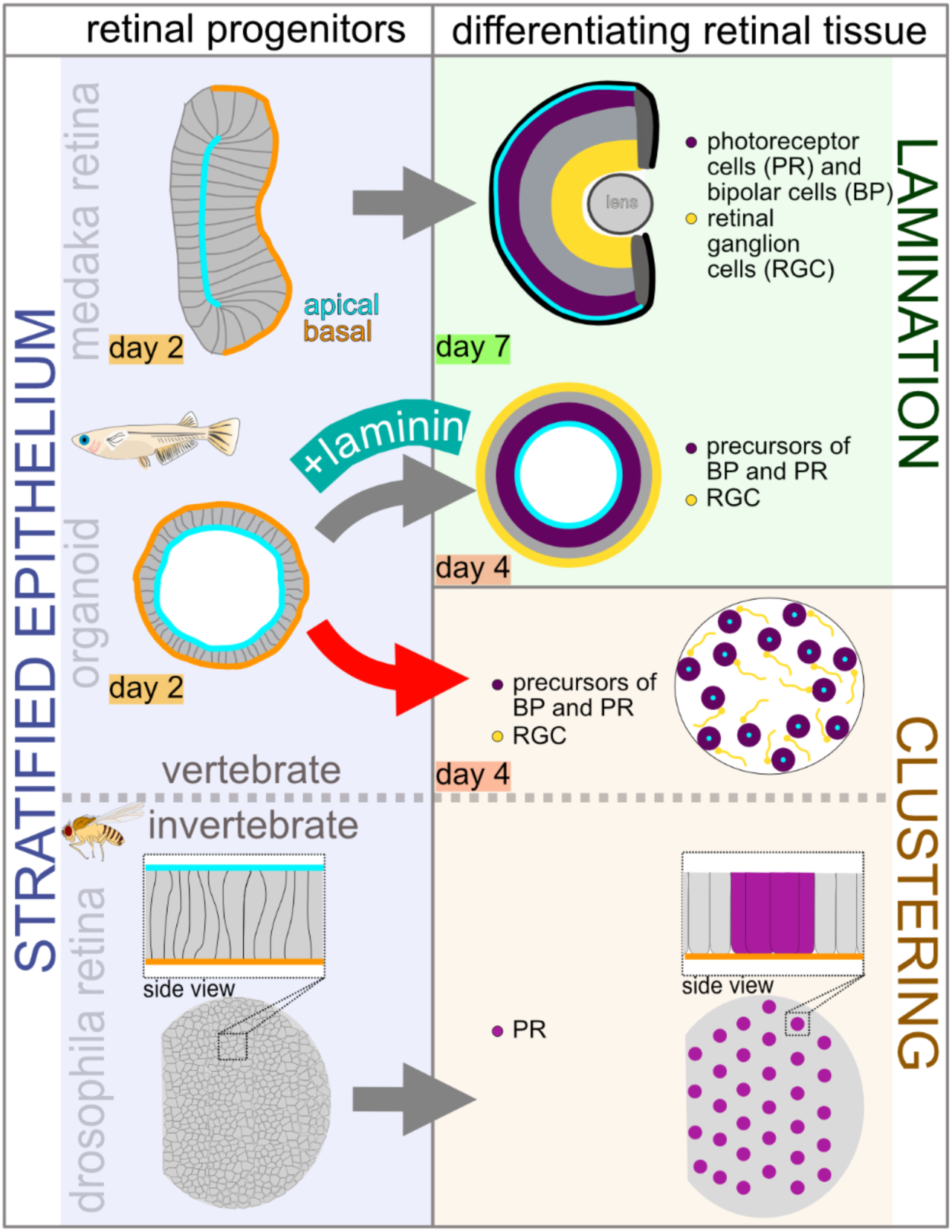

Retinal cells in medaka organoids adopt either a continuous layered epithelium when supported by laminin or a unit-based, ommatidia-like organization when epithelial continuity is lost. This dual outcome suggests that epithelial integrity represents a branching point between vertebrate and invertebrate strategies of retinal patterning, providing an experimental system to replay alternative evolutionary trajectories of eye design.

## 2. Results

### Retinal cell types in maturing retinal organoids emerge in spatially clustered domains

Medaka retinal organoids provide a rapid and tractable system to interrogate tissue patterning and polarity in a vertebrate context. Under previously established conditions (Zilova et al., 2021), aggregates of blastula-stage, pluripotent cells reproducibly adopt retinal identity already at day one of culture (Figure 1 A, B). Providing a basal polarity cue via Matrigel at this stage results in epithelialisation and the formation of a continuous layer of multipotent retinal progenitor cells (RPCs) by day two (Figure 1. A, B).

**Figure 1:**
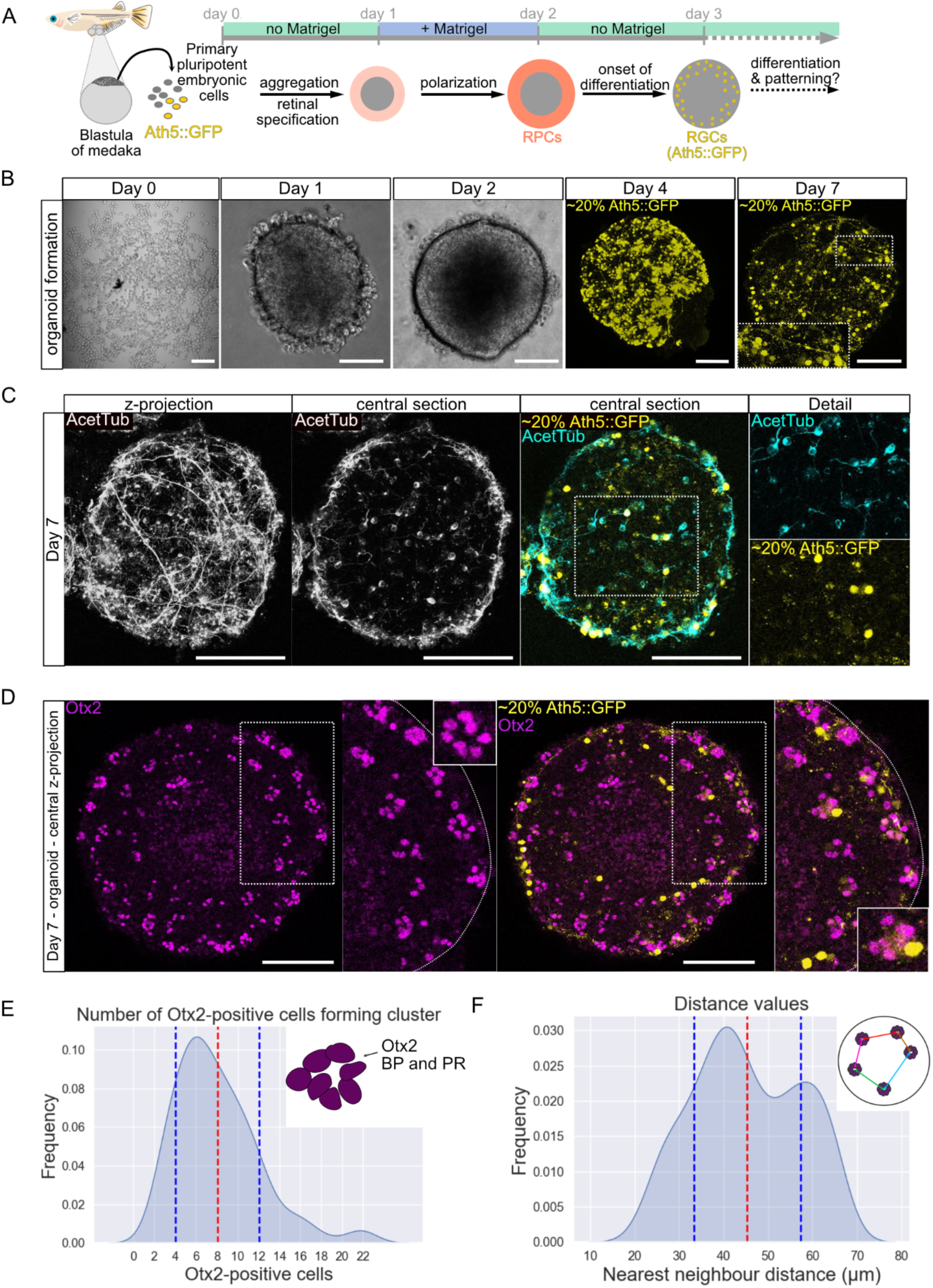
Neural retina in late medaka retinal organoids is structured in clusters. (A) Scheme of retinal tissue development in medaka retinal organoids as shown previously (Zilova et al., 2021). Neural retinal (NR) development of retinal organoids depicted from blastula stage (day 0) highlighting retinal tissue specification. Primary embryonic cells are isolated from blastula-staged medaka embryos and seeded for aggregation. Integrating cells from transgenic blastulas carrying a construct marking RGCs (Ath5::GFP) in the cell suspension (20%) enables tracing of RGCs. Aggregation and retinal specification are established by day 1 (red). Matrigel is added to the organoid culture on day 1, and by day 2, retinal progenitor cells (RPCs) are formed in the cortex of the organoid sphere (orange). More centrally (grey), the organoid contains non-retinal cells. Neural retinal cell differentiation is initiated by day 3 with the onset of RGC differentiation (yellow). (B) Representative images of organoids over the course of seven days. Single cells in culture wells on day zero (scale bar 200 μm) and the aggregated organoid on day one. By day four, RGCs (Ath5::GFP) expression indicates neuroretinal (NR) fate of the organoid. By day seven, RGCs are maintained and are visibly forming axonal projections throughout the organoid. Zoom on day seven organoid showing RGC with axon projecting centrally. Scale bar 100 μm for images day one to day seven. (C) Immunostaining for acetylated tubulin (AcetTub) marking axons in the retinal organoid at day seven. Axonal projections are visible in maximum intensity z-projection. Central section showing single neurons in overlay with RGC marker Ath5::GFP. Due to chimeric organoids carrying the Ath5::GFP label in ∼20% of RGCs, the acetylated tubulin label displays RGCs. (D) Otx2-positive cells (magenta) in the day seven retinal organoid form clusters in the cortex of the organoid. Maximum intensity z-projection of central slices in the organoid show distribution of Otx2-positive cells marking PR and BP and RGCs (Ath5::GFP). RGCs localize adjacent to Otx2-positive cell clusters. Scale bars 100 µm. (E) Quantification of overall cell number in Otx2-clusters in day seven organoids. Frequency of cell counts is plotted relative to Otx2-cell number counted (kernel density estimate (KDE) plot). The number of cells within one cluster ranges from 2-22 cells, the mean value is marked by red dashed line, standard deviation indicated by blue dashed lines. n=166 clusters were counted manually across n=18 organoids. (F) Distance between retinal cell clusters in direct neighborhood. Frequency of distance values between neighboring clusters plotted relative to the distance in μm (KDE plot). Mean distance to nearest neighbor is 45.34 μm (n=10 organoids from 2 independent experiments n=159 clusters, n=279 nearest neighbor distances).

From this organized neuroepithelium of RPCs, major neuroretinal cell types – including retinal ganglion cells (RGCs) and precursors of bipolar cells (BP), photoreceptors (PR), and horizontal cells (HC) (Zilova et al., 2021) – differentiate over the next days in the absence of any further exogenous growth factors or patterning cues (Figure 1 A, B).

To define the cellular repertoire and spatial arrangement of retinal cell types generated in late organoids, we analyzed day seven retinal organoids, corresponding to the developmental stage at which the embryonic medaka retina reaches full lamination (stage 39; Iwamatsu, 2004; Kitambi & Malicki, 2008).

We first focused on RGCs, the earliest-born retinal neurons (Hu & Easter, 1999; Kitambi & Malicki, 2008). Chimeric organoids generated by the co-aggregation of wild type cells with Ath5::GFP reporter cells (Del Bene et al., 2007; Zilova et al., 2021) labeled ∼20% of RGCs, facilitating the dynamic visualization of RGC emergence and distribution.

GFP-positive RGCs appeared consistently by day four (Figure 1B; Supplementary Figure 1), mirroring embryonic timing (Zilova et al., 2021; Del Bene et al., 2007; Kitambi & Malicki, 2008). Axonal projections were found from day four and maintained through day seven (Figure 1B; Supplementary Figure 1). Immunostaining of day seven organoids for acetylated tubulin highlighted axonal projections spanning both cortical and central regions of the organoid (Figure 1C). Overlay with Ath5::GFP confirmed label co-localization and indicated a broad distribution of RGCs throughout the organoid (Figure 1C). Together, these data establish that medaka retinal organoids faithfully recapitulate temporal neurogenesis and generate spatially distributed RGCs that extend axonal projections without external patterning inputs.

We next assessed the distribution of PR and BP cells by Otx2 immunostaining. Strikingly, Otx2-positive cells were not arranged in laminated layers in the cortex of the retinal organoid. Instead they were arranged into recurrent multicellular clusters, tightly associated with RGCs (Ath5::GFP) (Figure 1D). Quantitative analysis revealed that the clusters stereotypically contained eight cells on average (Figure 1E). Three-dimensional reconstruction further demonstrated that the clusters were evenly spaced within the cortical domain of the organoid, with nearest-neighbour distances displaying a bimodal distribution and an average spacing of ∼45 µm (Figure 1F). Altogether, these observations reveal that in the absence of extrinsic patterning and polarity cues, retinal cells in organoids self-organize into repeating, stereotypically sized and spaced clusters.

### Retinal clusters represent a structural unit with stereotypic composition

The observation that BP/PR co-positive cells occur in discrete clusters suggested that retinal neurons in organoids assemble in clustered, horizontal units rather than in the vertical, columnar arrangement, characteristic of the vertebrate retina. To determine whether these organoid-derived clusters represent a re-arranged version of the canonical retinal column, we assessed the cellular composition and 3D organization of cells within each unit.

To identify which retinal neurons differentiated in the organoids, we performed immunostaining for cell type-specific markers (Supplementary Figure 2). Otx2/Rx2 double-positive PRs were detected in every cluster (Figure 2A, B). We specifically identified both cone PR (Zpr1) and rod PR (rhodopsin) among the PR cells within the Otx2-clusters (Figure 2A, B). Horizontal cells (HC), characterized by the expression of Prox1, localize in close proximity to the cone PR marker (Figure 2A, B). BP cells were identified among the Otx2-positive cells by co-staining for PKCα, therefore localizing BP next to PR (Figure 2B). RGCs were consistently juxtaposed to the clusters (Figure 2A,C). Strikingly, we found that by day seven, all traceable neural retinal neurons (5 out of 6) differentiated in the organoids and were all either contained in or tightly associated with the Otx2-positive clusters described above (Figure 2A, B). Ultrastructural analysis by scanning electron microscopy confirmed tight cell–cell contacts within each cluster, as well as the presence of cilia, a hallmark of maturing PRs (Supplementary Figure 3, Supplementary movie 1) (Crespo and Knust, 2018, Malicki and Kitambi, 2008).

**Figure 2:**
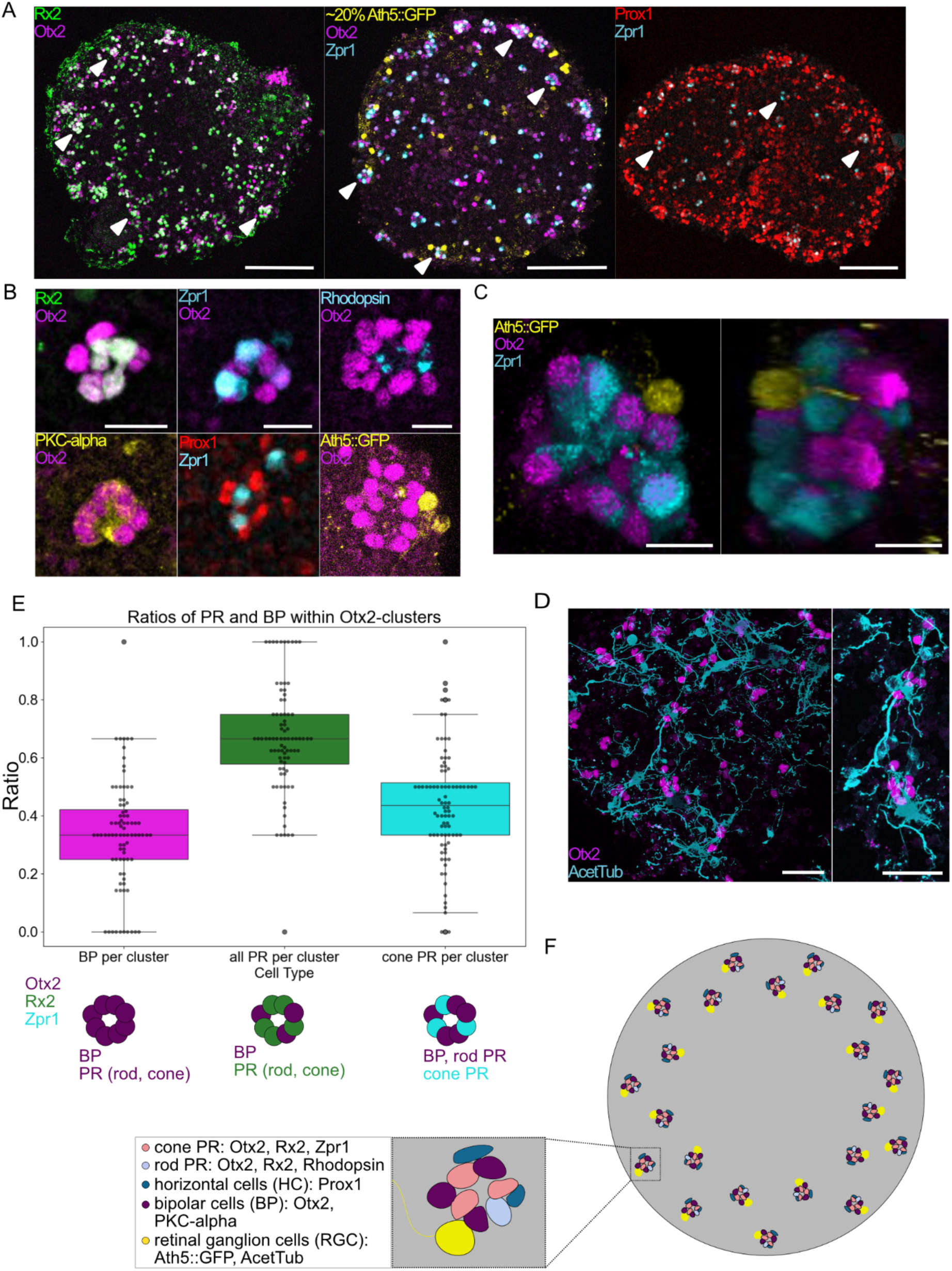
Retinal cell cluster include most neuroretinal cell types and form a recurrent structural unit. (A) Retinal cell marker for photoreceptors (PR) (Rx2), cone PR (Zpr1), horizontal cells (HC) (Prox1), retinal ganglion cells (RGCs) (Ath5::GFP) and the common marker for BP and PR (Otx2) across central sections of day seven organoids. Filled white arrow heads mark examples of clusters with cell types represented. Scale bars 100 µm. (B) Differentiated retinal cell types within and adjacent to Otx2-cell clusters. Otx2- and Rx2-double positive cells mark PR (n=16 organoids, in n=3 independent experiments) within cell clusters, and Otx2-positive Rx2-negative cells mark BP. PKC-alpha marking BP specifically (n=6 in n=2 independent experiments). Otx2- and Zpr1-positive cells represent cone PR (n=31 organoids, in n=6 independent experiments) and Otx2 - rhodopsin-positive cells mark rod PR (n=15 in n=3 independent experiments). Prox1-positive cells mark HC (n=16 organoids in n=4 independent experiments) adjacent to Zpr1-positive cone PR. Scale bar 10 µm. (C) 3D reconstruction of single cluster labelled for Otx2, Zpr1 and Ath5::GFP. 3D reconstructed cluster shown from two angles, rotated by 100°. Scale bar 10μm. (D) Immunostaining for Otx2 and acetylated tubulin markers shows the association of Otx2-labeled cell clusters with neurons. Axonal projections span between clusters. Scale bar 25 µm. (E) Comparison of relative contribution of BP, all PR and cone PR within cell clusters. Box plots showing ratios of numbers of BP per number of Otx2-positive cells per cluster (BP/Otx2-cluster) (median=0.33), all PR/Otx2 cluster (median=0.67) and cone PR/Otx2 cluster (median=0.44). Number of all PR quantified by manually counting the number of Rx2-Otx2-double positive cells within an Otx2-cell cluster and from the same data sets BP were scored as Otx2 only. Within n=9 organoids from 2 independent experiments, n=78 clusters were quantified. From different samples, Zpr1-Otx2-double positive cells were scored against Otx2-cell numbers within respective clusters. For n=9 organoids from 3 independent experiments, n=88 clusters were quantified manually. (F) Schematic summary of retinal cell types identified within retinal organoids at day seven, their respective labels and their positional relation.

In the vertebrate (fish) retina, PRs, BPs, and RGCs are arranged in distinct layers connected vertically via synaptic circuits to propagate visual information. In contrast, in organoids under the conditions described, these same cell types self-assembled into compact clusters, with PRs, BPs, and RGCs located in direct apposition (Figure 2A, B, C). The cellular composition was not only limited to a single cluster, but appeared recurrently across the organoid (Figure 2A). This was further validated by quantitative analysis, which revealed a stereotypic cellular composition. On average, Otx2-positive clusters consisted of ∼67% PRs and ∼33% BPs, with cones comprising two thirds of the PR population (∼44% of all Otx2-positive cells) (Figure 2D). This regularity was consistent across organoids, underscoring a recurrent “patterning logic” in which clusters combine the major cell types in reproducible ratios (Figure 2F). Complementary labeling with acetylated tubulin confirmed the regular apposition of these clusters to axon-bearing neurons, and tubulin-positive projections extended between clusters, hinting at emerging connectivity across units (Figure 2E).

These data establish that retinal cells in vertebrate organoids organize into discrete, reproducibly composed and distributed clusters that contain all elements establishing the retinal column as found in the maturing medaka retina at comparable stage. The emergence of such clusters stands in stark contrast to the canonical, layered architecture of vertebrate retinae.

### Reorganization of neural retinal tissue goes along with loss of epithelialization

During embryonic development of the medaka retina, proliferation, differentiation, and positioning of neuronal subtypes occur in a temporally ordered and spatially coordinated manner, emerging from a common pool of retinal progenitor cells (RPCs) (Kitambi and Malicki, 2008). Cellular ratios essential for retinal function are thought to be maintained either through lineage relationships or by signaling across neighboring progenitors (Avanesov & Malicki, 2010; Livesey and Cepko, 2011; Almeida et al., 2014; He et al., 2012). In later stages, studies on post-embryonic growth have shown that the full complement of retinal cell types arises from single stem cells, underscoring their intrinsic multipotency (Centanin, 2011).

To address the emergence of the retinal cell clusters, we assessed the origin of the cells within a cluster - either originating from a single or multiple progenitors. We performed clonal analysis in a chimeric assay by mixing blastula cells ubiquitously expressing GFP (20% EGFPubi) with unlabeled wild-type cells while seeding (Figure 3A). Lineage tracing revealed that all Otx2-positive clusters invariably contained both, labelled and unlabelled cells (Figure 3B). This indicates that clustering is not the result of a clonal expansion but rather reflects an active assembly. Thus, cells of different origin are assembling in the correct cell type ratios to form multicellular clusters.

**Figure 3:**
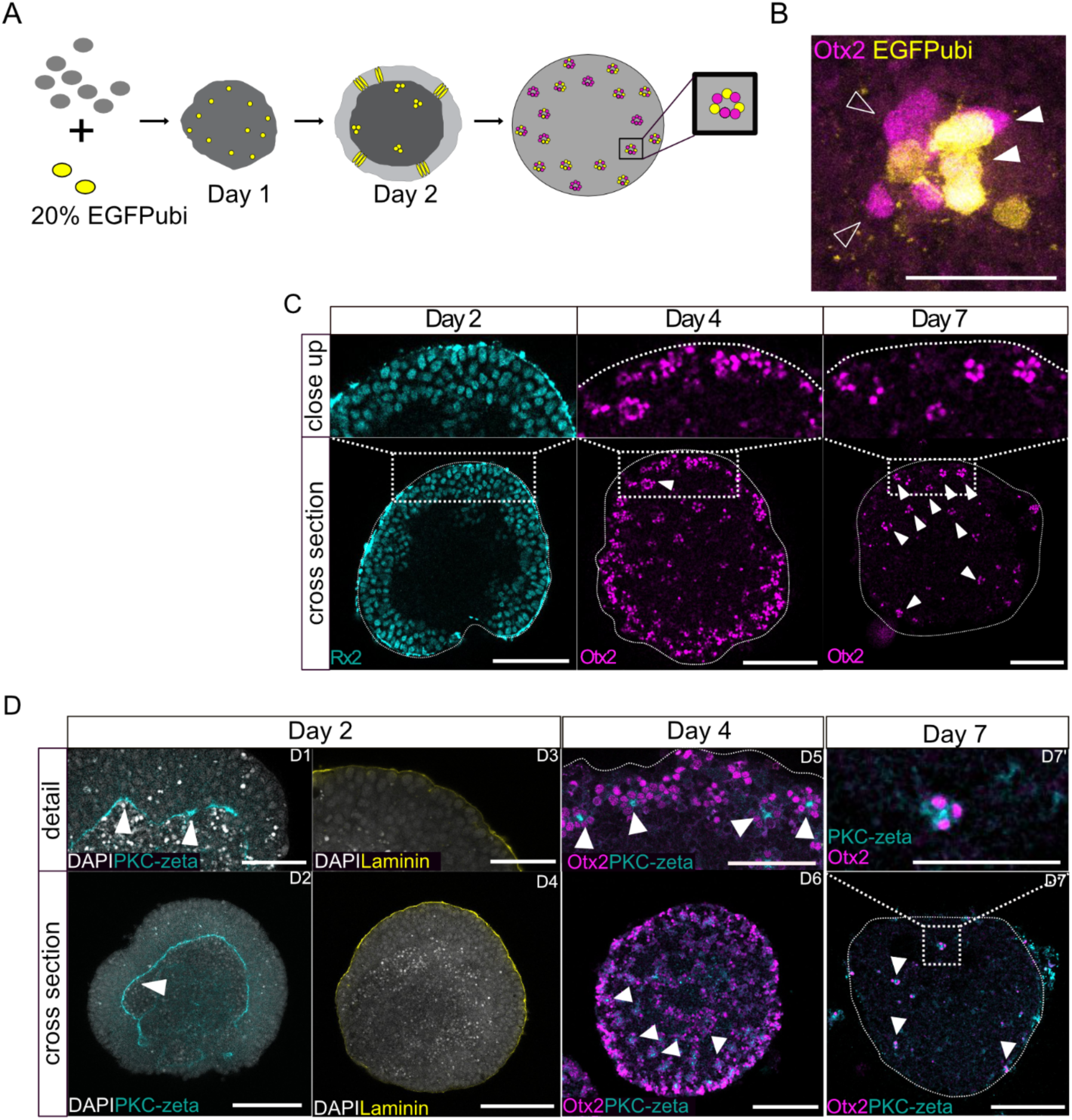
Progressive cluster formation along with loss of epithelial structure. (A) Scheme showing experimental set-up of clonal tracing in the retinal organoid and a schematic representation of the outcome. Unlabeled wild type cells are mixed with GFP-labelled cells while seeding organoids on day 0. Over the course of maturation, GFP-positive clones are expanding. Analyzing GFP-labeled and unlabeled cells within Otx2-positive clusters allows to follow up on clonal relationships of cells within a retinal cell cluster. (B) Otx2-expressing clusters are composed of cells of different clonal origin as seen in the mix of GFP-labelled cells (filled arrow head) and GFP-negative cells (empty arrow head). Scale bar 25 μm. (C) Retinal organoids stained for cellular distribution of retinal cells marked by Rx2 (RPCs) (day 2) or Otx2 on day four (n=100 organoids in n=14 independent experiments) and day seven (n=75 organoids in n=11 independent experiments). Otx2 expressing cell clusters are indicated by white filled arrowheads. Zoom on a central slice showing a cortical section in detail. Scale bar 100 μm. (D) Polarity of retinal tissue in retinal organoids from day 2 to day seven. Optical sections of organoids are shown. PKC-zeta marking apical side and Laminin marking basal side of cells. Polarity of RPCs on day 2 in retinal organoids show a global structuring as an epithelium with the apical (PKC-zeta) marker on inner surface of epithelium (marked by arrow head), and basal marker (Laminin) on the periphery. By day four, Otx2-positive clusters are formed in the organoid with an apical side towards the center of the cluster (marked by arrow heads). By day seven, the polarity is retained as the apical marker PKC-zeta can be found in the center of Otx2-cluster. The scale bar of organoid cross sections is 100 μm. The scale bar of organoid close ups is 50 μm.

We next followed the formation of clusters over time to define a temporal window relevant for the structural organization of the tissue. On day two, RPCs established a continuous epithelial sheet of cells organized in vertical orientation relative to the cortex plane. By day four, neural retinal cell differentiation was initiated with Otx2-positive PR and BP precursors emerging and distributed predominantly at the organoid cortex (Figure 3C) (Zilova et al., 2021). At this stage, first indications of Otx2-positive cell clustering became apparent (Figure 3C). By day seven, all Otx2-positive cells had organised into discrete clusters, spaced at regular intervals across the organoid cortex (Figure 3C). Notably, the clustered phenotype persisted stably beyond day seven (day nine and day eleven; Supplementary Figure 4), consistent with the establishment of a constructive and self-maintained architecture.

Cellular positioning of the differentiating cell types in the embryonic retina is oriented within the apico-basal axis of the neuroepithelium. The clustered organization in the retinal organoid suggests that the epithelial organization is affected. We therefore traced epithelial polarity over time by analyzing the distribution of apical and basal polarity markers in the retinal organoid tissue. We found that on day two, RPCs exhibited a coherent epithelial organization with apical domains facing the organoid core and basal domains exposed at the surface, a pattern comparable to the optic cup *in vivo* (Figure 3D, Supplementary Figure 5). However, between day two and day four, this global epithelial organization is progressively lost as seen by the distribution of differentiating retinal cells marked by Otx2. The emergence of Otx2-positive cell clusters is occurring while the common apical side of the former epithelium is not detectable anymore by the apical polarity marker (PKC-zeta). By day seven, the polarity of the tissue was reorganized to a local pattern: each cluster established its own apico–basal orientation, with apical domains directed toward the cluster center (Figure 3D). This local polarity pattern remained stable through subsequent stages. In contrast, the embryonic medaka retina maintains a continuous apico–basal polarity throughout development, with PRs localized apically and RGCs basally (Supplementary Figure 5).

Thus, the transition from a continuous epithelium to discrete retinal clusters represents a structural re-organization unique to the *in vitro* environment and not observed in vertebrate retinal development. The loss of tissue-wide epithelialization suggests that external inputs—present in the embryonic context but absent in the artificial culture environment—are required to maintain the continuous epithelial organization necessary for layered retinal architecture.

### Laminin supports epithelial structure in retinal organoids

The establishment and maintenance of epithelial organization in the retina is closely linked to the extracellular matrix (ECM), with laminins representing a key component of retinal ECM (reviewed in Prieto-Lopez at al., 2024; Dorgau et al., 2018). In our organoid system from day two onwards, cultures are maintained without ECM supplementation. To test whether external cues, such as laminin, are sufficient to preserve epithelial tissue architecture during the onset of retinal differentiation, we supplemented cultures with laminin starting at day two. This time point is prior to the breakdown of epithelial layering observed in standard conditions (Figure 4A). Indeed, organoids retained integrity of epithelia and layering of retinal cell types by day four when cultured with external addition of laminin (Figure 4B, Supplementary Figure 6). Importantly, tissue polarity was preserved, with the apical side of the epithelium oriented toward the organoid core. Thus, laminin provides the external structural cue that is absent in standard culture conditions and is sufficient to maintain epithelial tissue continuity. Within these organoids, Otx2-positive cells localized apically, while RGCs occupied the outer cortex, mirroring the relative horizontal arrangement of cell types in the embryonic retina.

**Figure 4:**
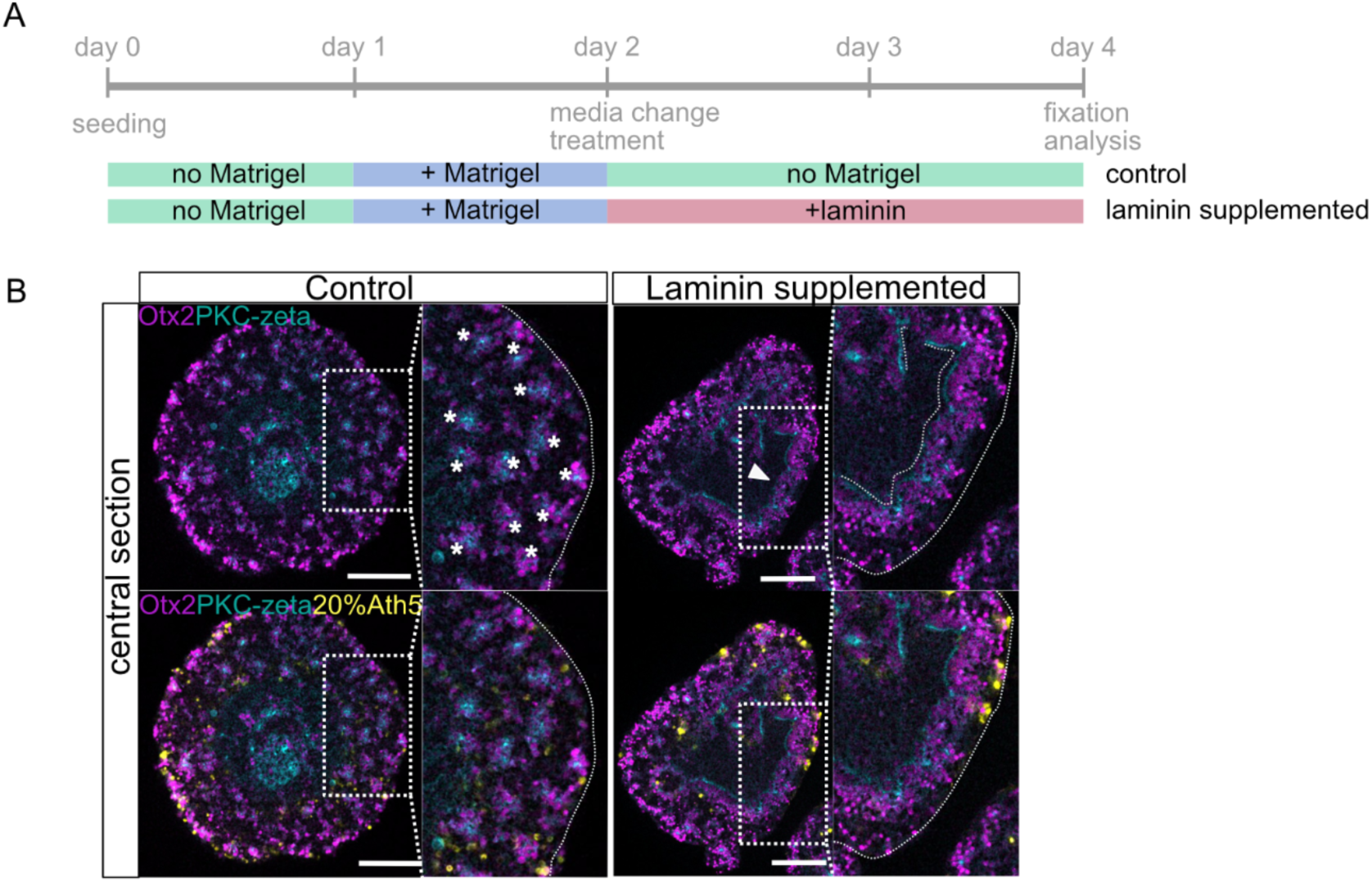
Supplementation of organoids with Laminin is sufficient to restore epithelial layering of neurons in retinal organoids. (A) Experimental outline of organoid culture supplemented with 25 μg/ml Laminin from day two to day four. Organoids were grown until late day two under standard conditions (control) and addition of 2% Matrigel on day 1. Preceding the loss of tissue polarity, media was changed and for Laminin-exposed samples, 25 μg/ml Laminin was added to the well. Samples were incubated until day four. Afterwards the samples were fixed, immunostained and imaged. (B) Loss of layering upon progression of organoid development after day 2 can be partially prevented by Laminin-supplementation by day 2. In Laminin-supplemented culture, tissue stretches stay polarized by day four and cluster formation is prevented. Otx2-positive cells line up with apical pole towards the center and RGCs arrange in a layer adjacent facing the outer rim of the organoid. In five independent experiments, with a total number of n=28 organoids, n=24 (85%) showed tissue polarization after laminin-treatment. Scale bar 100 μm.

The preservation of epithelial layering and the pseudostratified organization of retinal cell types as in the fish retina by the presence of laminin underscores the critical role of ECM in instructing epithelial tissue architecture and polarity. In contrast, organoids grown without laminin abandon epithelial layering and instead adopt a clustered organization of retinal cells. The process of epithelial destruction and retinal cell clustering seen in the retinal organoids marks a striking alternative retinal tissue architecture for a vertebrate retinal model.

## 3. Discussion

Our study identifies laminin as a critical extracellular cue for maintaining epithelial organization in medaka retinal organoids.

When laminin is not provided exogenously, epithelial continuity is lost and retinal cells adopt a clustered, unit-based arrangement. Laminin supplementation preserves epithelial layering and polarity, resulting in a layered arrangement with Otx2-positive cells apically and RGCs basally, thereby resembling the vertical organization of a retinal column in the embryonic retina. These findings highlight the instructive role of ECM in organizing retinal tissue architecture and uncover an alternative patterning mode that emerges once epithelial integrity is not sustained.

Other vertebrate retinal organoid models based on mouse or human cells achieve layered tissue organization without prolonged exogenous ECM exposure, suggesting that mammalian retinal cells provide laminin and other ECM components autonomously (Eiraku et al., 2011; Nakano et al., 2012; Dorgau et al., 2018). In medaka retinal organoids, however, we showed that the laminar structuring of the retinal cell types depends on exogenous supplementation of laminin. This species-specific difference in self-sufficiency highlights how variations in ECM contribution bias tissue architecture toward layered versus modular outcomes.

The clustered assemblies formed in laminin-free cultures suggest that, in the absence of epithelial organizing signals, retinal cells follow an intrinsically encoded or physically advantageous strategy. RGCs, the first born retinal cell type, could form a nucleation point for the forming retinal cell clusters (Icha et al., 2016). The recurrent inclusion of PRs, BPs adjacent to RGCs within these clusters points to a regulated process faithfully assembling the essential components of the retinal column. This unit-based organization is strikingly reminiscent of ommatidial patterning in the *Drosophila* retina, where photoreceptor clusters arise as structurally and functionally independent modules. Ommatidia emerge from an initially polarized epithelium, with unit formation proceeding concomitantly with the establishment of the morphogenetic furrow and the perpendicular patterning axis (Warren & Kumar, 2023). The striking resemblance of laminin-free retinal organoids to the unit-based architecture of the invertebrate eye suggests that the breakdown of epithelial organization may underlie divergent patterning strategies of retinae across the animal kingdom. Our observations therefore support the view that epithelial continuity represents a key decision point between layered and unit-based architectures of the retina.

Taken together, our findings raise the possibility that the evolutionary divergence of vertebrate and invertebrate eye types reflects alternative solutions to the challenge of organizing photoreceptor circuits: continuous epithelial layering, supported by ECM, versus modular unit formation upon epithelial modulation. The medaka organoid system, in which both modes can be elicited, thus provides the unique opportunity to dissect polarity-dependent retinal structuring and to experimentally explore potential evolutionary trajectories in eye design.

More generally, organoid systems reveal latent self-organizing capacities that are normally masked by embryonic constraints. By uncovering alternative tissue architectures, such as layered epithelia versus unit-based modules, organoids not only serve as models for development and disease but also as experimental windows into evolutionary processes and the fundamental principles of tissue patterning.

In this sense, retinal organoids offer a rare experimental system to replay alternative evolutionary scenarios of eye design in real time.

## 4. Supplement

**Supplement Figure 1 (related to Figure 1):**
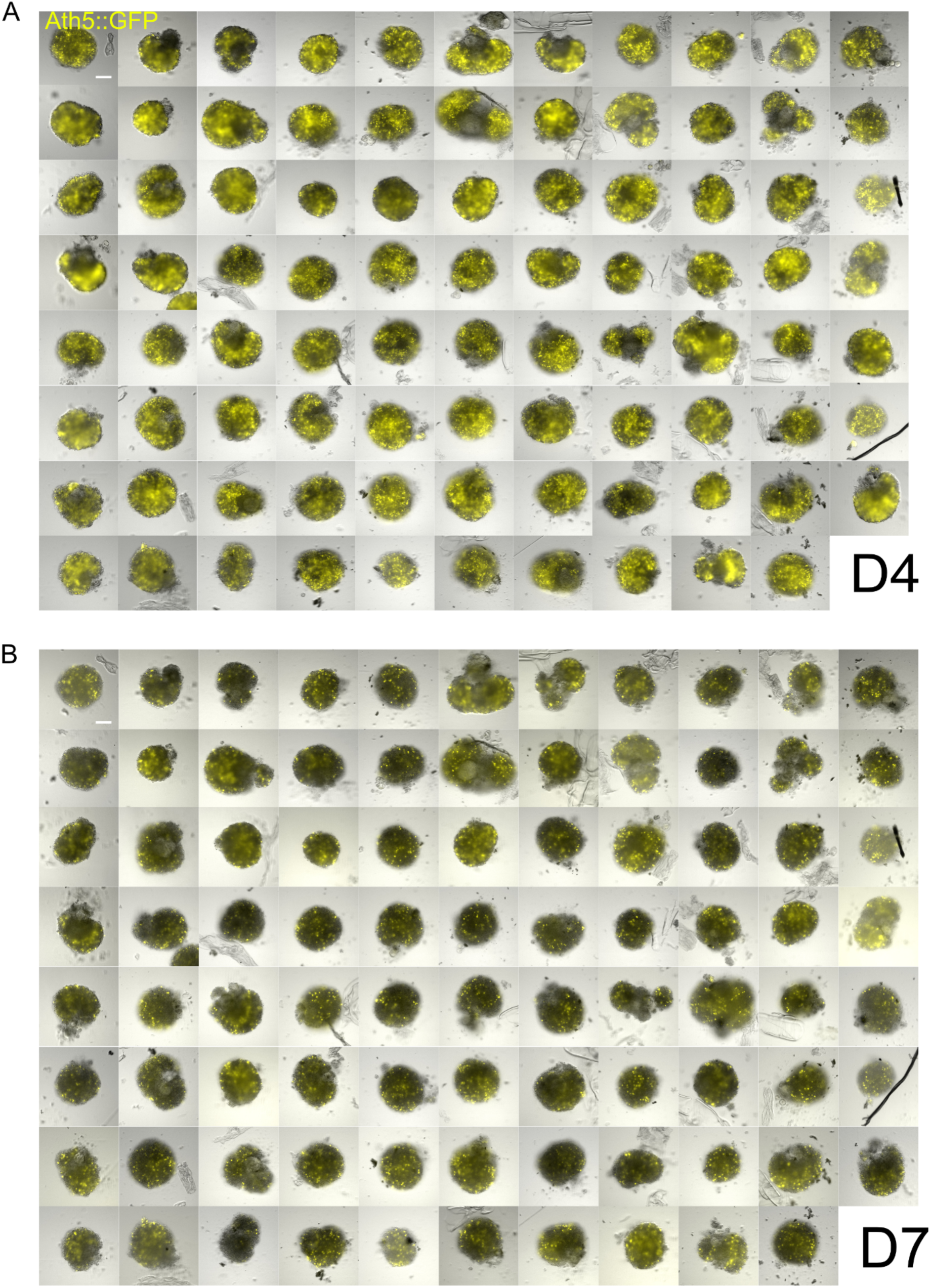
Differentiation of retinal organoids containing RGCs is robust and reproducible. Retinal organoids grown on one 96 well plate imaged with the Acquifer imaging machine on day four (A) and day seven (B). GFP signal of Ath5::GFP reporter shown in overlay with brightfield images. Scale bar 100 µm.

**Supplement Figure 2 (related to Figure 2):**
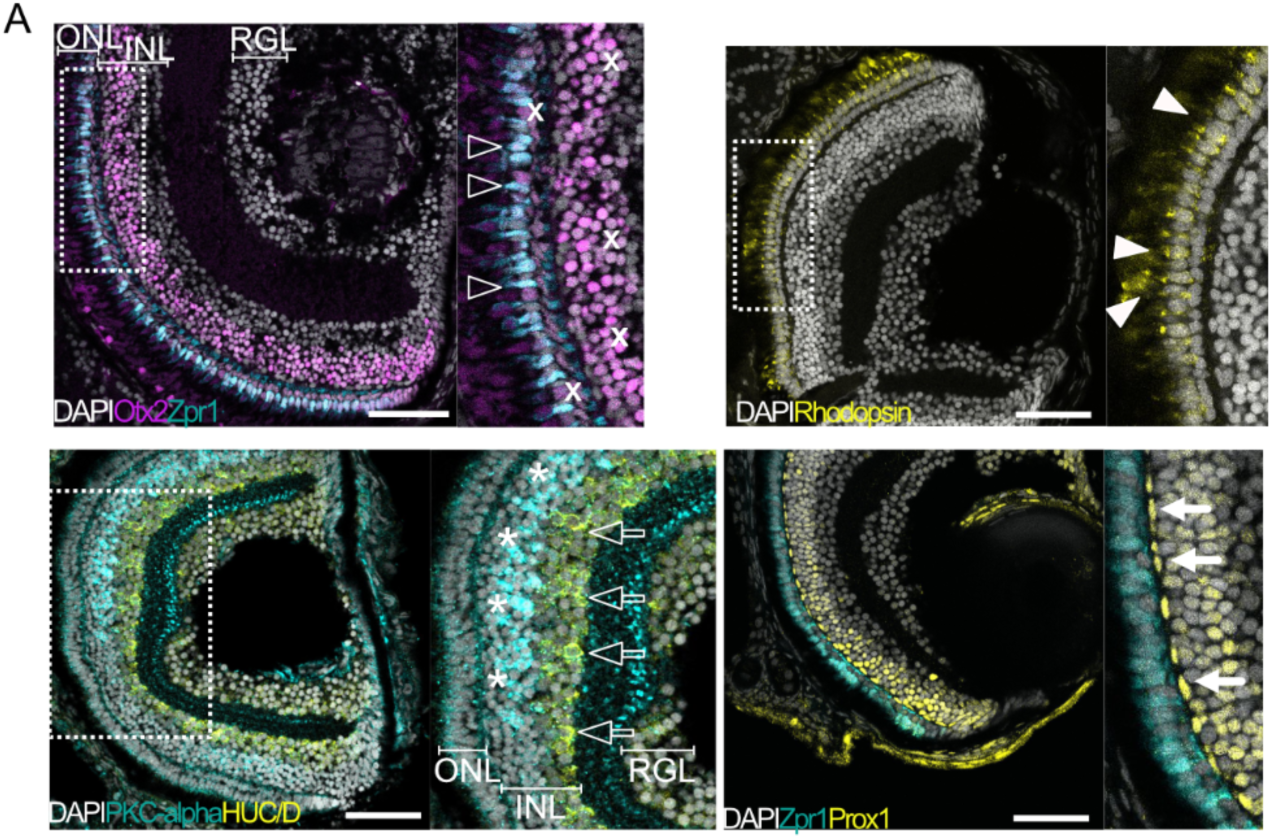
Retinal cell type distribution in embryonic retina. Retinal cell types labeled with cell type-specific antibodies in sectioned fish retina at day 8 (0 days post hatch). Otx2 labels photoreceptors (PRs) and bipolar cells (BPs), Zpr1 and Otx2 double labeling showing cone PR. Rhodopsin marking rod PR. Prox1 marking horizontal cells adjacent to Zpr1-positive cone PR. PKC-alpha marking BP and HuC/D marking amacrine cells and retinal ganglion cells (RGCs). Scale bar 50 μm.

**Supplement Figure 3 (related to Figure 2):**
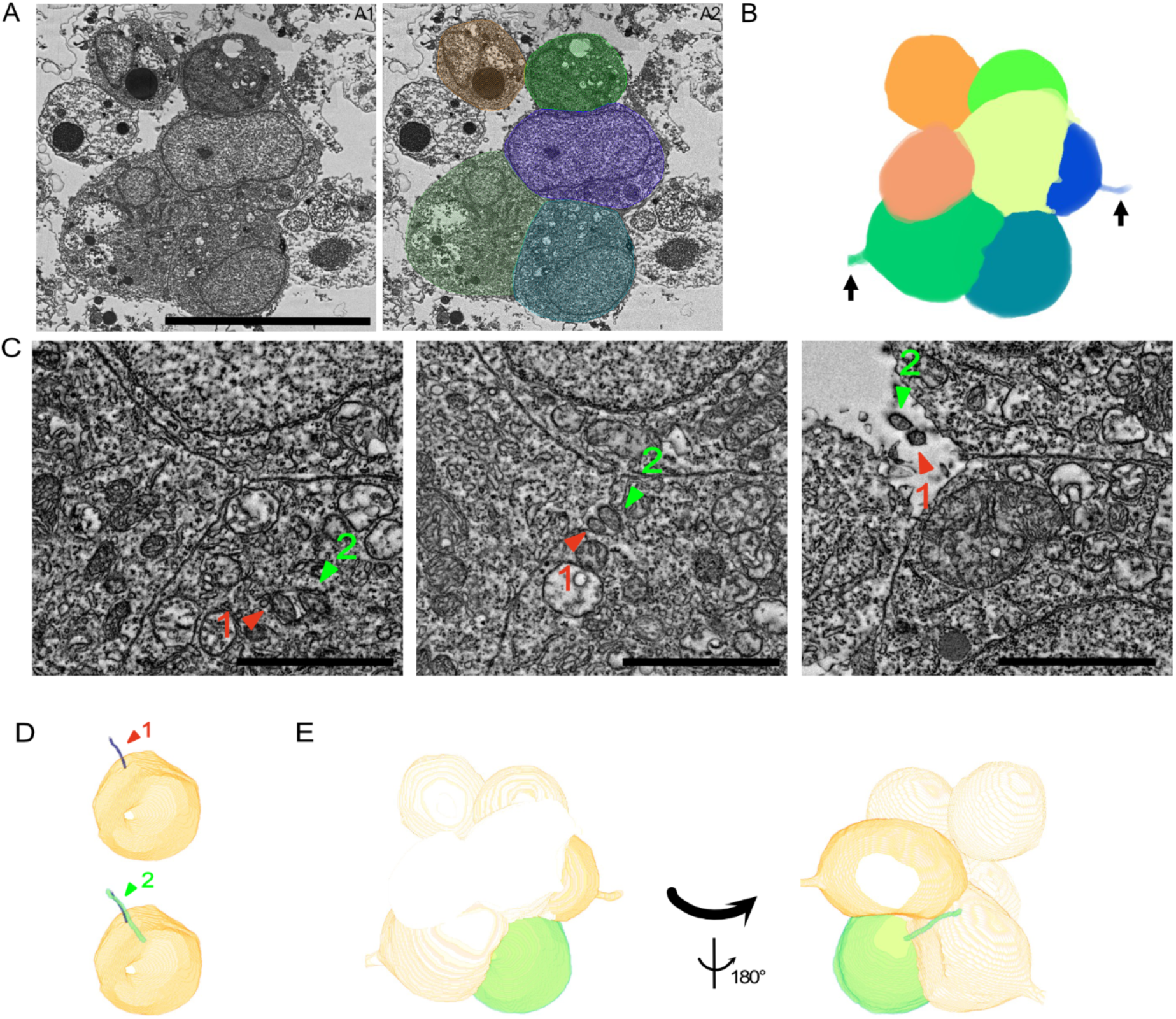
Ultrastructure of retinal cell clusters show tight association of cells and cilia (A) Example of cell cluster in day seven organoid imaged with electron microscopy. Single z-plane of cell cluster showing 5 of 7 tightly associated cells within the cluster. The same plane is overlayed with segmentation markup using Amira. Scale bar 10μm. (B) 3D projection of seven segmented cells forming a cluster. Three cells show axonal projections. Two of those are visible in the displayed orientation (indicated by arrows). (C) Cell with two cilia projecting into the same direction and out of cell. Cilia grow out along neighboring cells. Cilia marked by arrows and numbers. Scale bar 2μm. (D) 3D volume of segmented cell in (C) with cilium1 only (top) and both cilia (bottom). (E) Cell with cilia (green) in context with whole cell cluster shown in cell segmentation. Shown in same orientation as single cell display in (D) and in context as in (B) and turned by 180° on the vertical axis.

Supplemental movie 1: Extending cilia from cell within the retinal cluster visualized with electron microscopy. Cilium 1 (red) and cilium 2 (green) outlined in image stack. Scale bar 5µm.

**Supplement Figure 4 (related to Figure 3):**
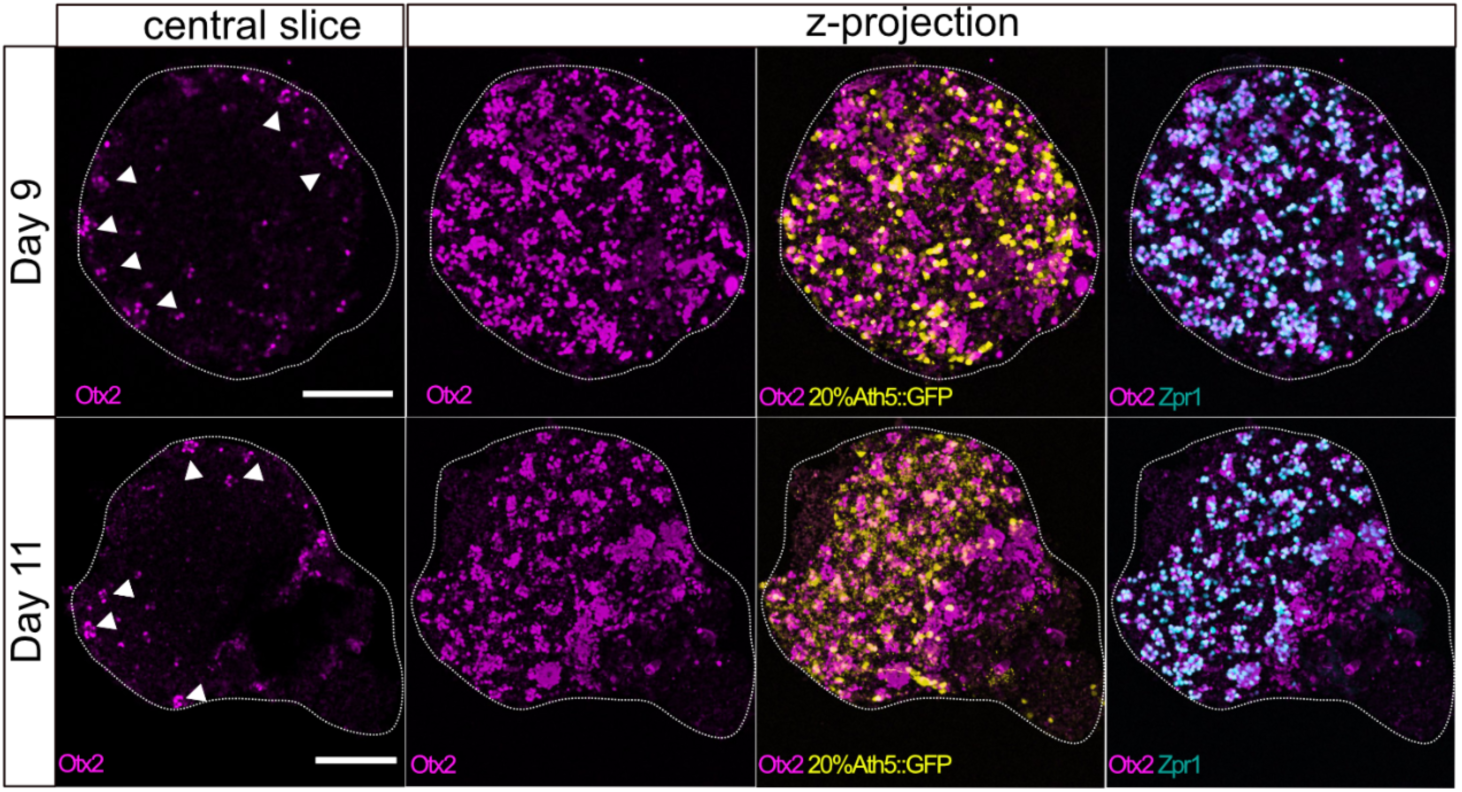
Retinal clusters are stable until at least day eleven. Retinal organoids derived of wild type (wt) and Ath5::GFP reporter cells (4:1) stained for the presence of Otx2, Zpr1 and GFP by immunohistochemistry on day 9 (n=11 organoids in N=3 independent experiments) and day eleven (n=15 organoids in n=3 independent experiments). Otx2 expressing cell clusters are indicated by white arrow heads. Scale bar 100 μm.

**Supplement Figure 5 (related to Figure 3):**
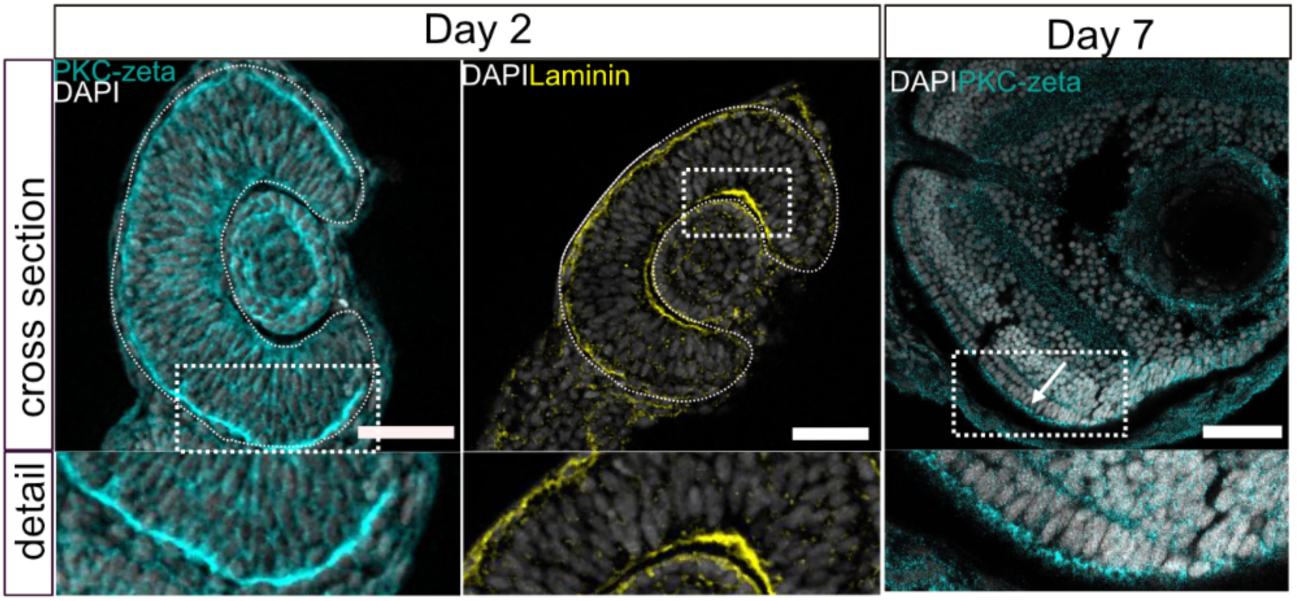
Fish retina polarity Polarity of retinal tissue in medaka embryos on day 2 and day seven. Optical cross sections of embryonic retinae are shown. PKC-zeta marking apical cell polarity and Laminin marking basal pole of cells. Apical pole of the retinal tissue within the optic cup is facing the brain, while the basal side of the tissue is facing the lens. At day seven, the apical polarity in PR is still present at the interface to the RPE. Scale bar 50μm.

**Supplement Figure 6 (related to Figure 4):**
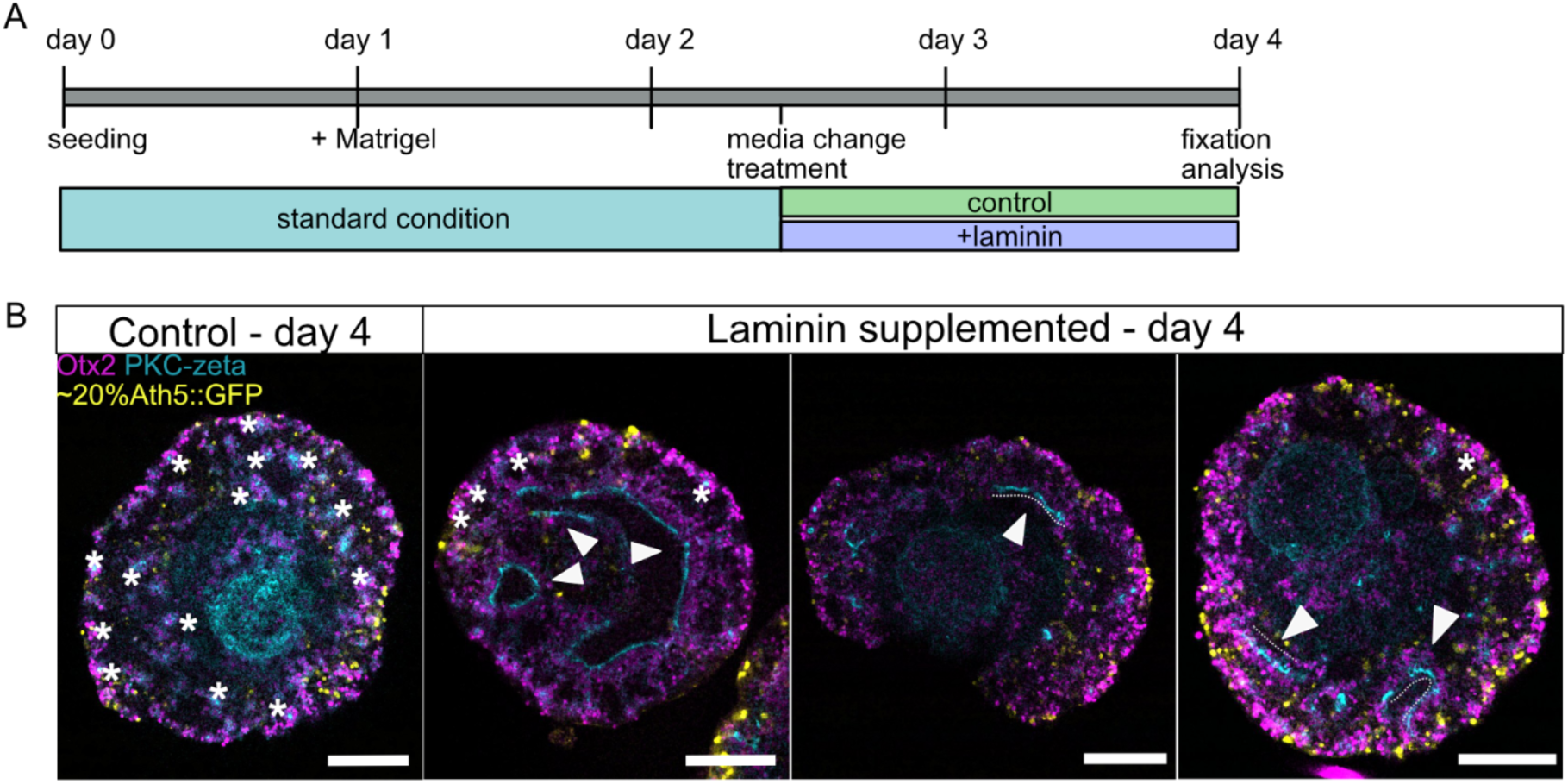
Phenotypic spectrum of organoids grown in laminin-supplemented culture. (A) Experimental procedure for laminin-supplemented organoid culture. After culturing organoids in standard conditions until late day 2, laminin was added to the ‘treated’ organoid. Samples were cultured until day four and then processed for antibody staining. (B) Tissue structure in standard and laminin-supplemented culture. By day four, tissue stretches remain polarized visible by Otx2-positive cells (magenta) lining up with the apical side (PKC-zeta, cyan) towards the core of the organoid (marked by arrows). Clusters can be found in other areas of the laminin-enriched organoids (marked by stars). Control organoids exclusively show clustered phenotype (marked by stars). Ath5::GFP marks fraction (∼20%) of RGCs and their distribution. Scale bar 100 μm.

## 5. Methods

### Fish husbandry

Medaka (*Oryzias latipes*) stocks were maintained at 28°C in closed stocks with a 14 h light and 10 h dark cycle. Husbandry of the fish (permit number: AZ35-9185.64/BH; line generation permit: 35–9185.81/G-145/15 Wittbrodt) was performed according to the EU directive 2010/63/EU guidelines and the German animal welfare laws (Tierschutzgesetz §11, Abs. 1, Nr.1). Fish lines used were Cab (Loosli et al., 2000) as wild type, Ath5::GFP (Del Bene et al., 2007) and Wimbledon dsTrap#6 line (Centanin et al., 2011) here called EGFPubi.

### Medaka retinal organoid generation

Medaka organoids were prepared from late blastula stage primary embryonic stem cells as described in Zilova et al. (2021). Cells were extracted from stage 10 (Iwamatsu, 2004) by dechorionating embryos using hatching enzyme. The cell mass was manually detached from the yolk, washed and dissociated in 1x PBS (Thermo Scientific, Cat# 100110023). The cell suspension was strained by centrifugation for 3 min at 180*g and re-suspended in ‘differentiation medium’ (DMEM/F12 (Gibco, Cat# 21041025), 5% knockout serum replacement (KSR) (Gibco, Cat# 10828028), 0.1mM MEM non-essential amino acids (Sima Aldrich, Cat# M7145), 0.1 mM sodium pyruvate (Sigma Aldrich, Cat# S8636), 0.1 mM β-mercaptoethanol (Gibco, Cat# 21985023), 2mM HEPES pH 7.4 (Carl Roth, Cat# 7365-45-9), 100 U ml-1 penicillin-streptomycin (Gibco, Cat# 15140122)). Per single organoid, 1500 cells in 100 µl were seeded into individual wells of a low binding U-bottom-shaped 96-well plate (Nunclon Sphera U-Shaped Bottom Microplate, Thermo Fisher Scientific, Cat# 174925). Cells of different genetic backgrounds were combined when indicated (20% Ath5::GFP with 80% Cab cells; 20% dsTrap#6/ubi::GFP with 80% Cab cells). The culture was incubated at 26°C without CO_2_ control over night (o.n.) and organoids were washed in differentiation medium early day 1. Under the standard conditions, day 1 organoids were supplemented with Matrigel (Corning, Cat# 356230) to a final concentration of 2%. From that timepoint on, organoids were incubated at 26°C with 5% CO_2_. On day 2, organoids were transferred into ‘maturation medium’ (DMEM/F12 (Gibco, Cat# 21041025), 10% fetal bovine serum (Sigma Aldrich, Cat# 12103C), 1x N2 supplement (Gibco, Cat# 17502048), 1mM taurine (Sigma Aldrich,Cat# T8691), 20mM HEPES pH 7.4 (Carl Roth, Cat# 7365-45-9), 100 U ml-1 penicillin streptomycin (Giboc, Cat# 15140122)) and media was changed on day 4 and day 7 for fresh maturation media.

For supplementation of organoids with laminin from day 2, laminin protein solution (Roche, Cat# 11243217001) was added to a final concentration of 25 µg/ml to one organoid per well. For the control condition, standard culture was performed as described above. Organoids were incubated at 26°C with 5% CO_2_ control until day 4.

### Whole mount staining of medaka organoids

Organoids were fixed in 4% PFA (paraformaldehyde) (Sigma Aldrich, Cat# P46148) in 1x PTW (1x PBS, 0.05% Tween20) for 48h at 4°C. Samples were washed with 1x PTW and permeabilized with Aceton (Sigma Aldrich, Cat# 32201-2.5l) at −20°C for 15 min. Organoids were washed in 1xPTW and blocked in 10% BSA (Sigma Aldrich, Cat# A9418) in 1xPTW for 2h at r.t.. Primary antibodies (ABs) (mouse anti-acetylated tubulin (Sigma Aldrich, Cat# mA1-12717); chicken anti-GFP (Thermo Fisher Scientific, Cat# A10262); rabbit anti-Laminin (Sigma-Aldrich, Cat# L9393); goat anti-Otx2 (R&D systems, Cat# AF1979); rabbit anti-PKC-alpha C-20 (Santa Cruz, Cat# sc-208); rabbit anti-PKC-zeta C-20 (Santa Cruz, Cat# sc-216); rabbit anti-Prox1 (Millipore, Cat# AB5475); mouse anti-Rhodopsin (Sigma-Aldrich, Cat# MABN15); rabbit anti-Rx2 (Reinhardt et al., 2015); mouse anti-Zpr1 (Zebrafish International Resource Center)) were applied in 1% BSA in 1x PTW and incubated for 48h at 4°C. Samples were washed in 1x PTW and the secondary ABs (Invitrogen) and DAPI (1:500) were applied in 1% BSA in 1x PTW o.n. at 4°C in darkness. Afterwards, samples were washed in 1x PTW. For confocal microscopy, organoids were mounted in optical clearing solution (Zhu et al., 2019).

### Cryosectioning and immune staining of medaka hatchlings

Medaka hatchlings (0 days post hatch) were anaesthesized and fixed in 4% PFA in 1xPTW for 48h at 4°C. Heads were manually dissected using forceps. Samples were cryopreserved in 30% (w/w) sucrose and sectioned to 16 μm sections in Tissue Freezing Medium (Leica, Cat# 1402018926) Sections were rehydrated with 1x PTW for 10 min and blocked with 10% BSA in 1x PTW for 2h in a humidified chamber. After washing in 1x PTW, primary ABs (anti-GFP chicken (Thermo Fisher Scientific, Cat# A10262); anti-Laminin rabbit (Sigma-Aldrich, Cat# L9393); anti-Otx2 goat (R&D systems, Cat# AF1979); anti-PKC-alpha C-20 rabbit (Santa Cruz, Cat# sc-208); anti-PKC-zeta C-20 rabbit (Santa Cruz, Cat# sc-216); anti-Prox1 rabbit (Millipore, Cat# AB5475); anti-Rhodopsin mouse (Sigma-Aldrich, Cat# MABN15); anti-Rx2 rabbit (Reinhardt et al., 2015); anti-Zpr1 mouse (Zebrafish International Resource Center)) were applied in 1% BSA o/n at 4°C in a humidified chamber. Sections were washed in 1x PTW and secondary ABs (Invitrogen) and DAPI (1:500) were applied in 1% BSA in 1x PTW and incubated for 2 h at 37°C in a humidified chamber. Afterwards, samples were washed in 1x PTW and mounted by adding 60% glycerol (Sigma Aldrich, Cat# V900122).

### Light microscopy imaging and image processing

Live imaging of organoids was done using a ACQUIFER (ACQUIFER Imaging GmbH) imaging machine. Fixed organoid and embryo samples were imaged using a Leica TCS Sp8 Dmi8 inverted confocal microscope (20x and 63x oil immersion objective). For display, confocal images were subjected to rolling ball background subtraction of 25 or 50 pixels and median filtering (1.2 pixels) using the Fiji software. Within one condition, all images were treated the same way.

Cell type ratios and cell numbers within late retinal organoids were quantified manually using Fiji. The number of cells within a cluster was counted by using the cell counter tool. A cluster was defined as a group of Otx2-positive cells with two or more cells, which are located in close vicinity. Counting cell types within clusters was done using the cell counter tool and scoring the double positive cells for the respective cell type specific label (Rx2-Otx2 – PR; Zpr1-Otx2 – cone PR) within an Otx2-positive cell cluster. Otx2-only cells in the Rx2-Otx2-double labeling were counted as BP cells. Distance measurements between Otx2-clusters was done by defining the center of each cluster in a defined z-stack (sub-stack; up to 30 clusters per stack) and documenting the position in 3D by recording the coordinates (measuring tool) using Fiji (ImageJ2). To define the distance between all clusters of a sub-stack, the vector length between all coordinates within one sub-stack were calculated (n=159 clusters in n=10 organoids; 2-3 sub-stacks per organoid). The data sets (‘all distances’) include the distances between not directly neighbouring clusters, which were removed as outliers. To filter for the distances of direct neighbouring clusters only (remove outliers), a range of values was defined by the average and 3x standard variation (+/-) of the smallest distance value within the distance data of each sub-stack. This range was used as boundaries within the ‘all distances’ data and the values were displayed using a kde (kernel density estimate) plot. Graphs were plotted and statistical analysis done in Python (Jupyter lab interface).

### Electron microscopy Sample Preparation and Ultramicrotomy

Organoids (day seven) were fixed in 0.1 M PIPES buffer (pH 6.8.) (Carl Roth, Cat# 9156.3) + 1.25% glutaraldehyde (Plano, Cat# R1010) + 2% PFA (Plano, Cat# R1026) for several days. After washing with 0.1 M PIPES, the samples were incubated in 1% osmium tetroxide (OsO_4_) (Science Services, Cat# E19152) and 0.8% potassium ferricyanide (K₃[Fe(CN)₆]) (Science Services) for 1 hour on ice. Subsequently, the samples were stained overnight with 2% uranyl acetate (Science Services, Cat# E22400) in 25% ethanol/water.

Samples were then subjected to a graded ethanol dehydration series (25%, 50%, 70%, 90%, and 100% ethanol/water), followed by ethanol/acetone (1:1) and 100% acetone for 15 minutes/dilution step. Samples were then infiltrated with Epon resin (42.4 g glycid ether 100 (Serva, Cat# 21045.01), 29.6 g dodecenylsuccinic acid anhydride (DDSA) (Serva, Cat# 20755.01), 18.4 g methyl nadic anhydride (MNA) (Serva, Cat# 29452.01), 2.4 g benzyldimethylamine (BDMA) (Serva) as initiator) first with 30% acetone, then with 70% Epon in acetone, each step for 2 hours. Embedding of samples was done in 100% Epon resin. Polymerization was done at 60°C for 2 days.

Serial sections (100 nm thickness) were prepared using a PowerTome Ultramicrotome (RMC Boeckeler) (Jumbo 35° diamond knife (Diatome, Switzerland)). For further analysis, sections were transferred onto silicon wafer substrates. For post-staining the substrates (with sections facing down), the sections were placed on a 200 µl droplet of 3% uranyl acetate (Science Services, Cat# E22400) in water for 10 minutes, washed with water, and then immersed in a solution of 3% lead citrate (Science Services, Cat# DM22410) in water for 5 minutes.

### Electron Microscopy and Image Processing

Sections were imaged using a field emission scanning electron microscope (Ultra 55, Carl Zeiss Microscopy) operated at a primary electron energy of 1.5 keV. Secondary electron (SE2) and backscattered electron (ESB) detectors were used to capture images. Using the Atlas 5 software (Carl Zeiss Microscopy), automated acquisition of large scan fields was performed. The resulting image stacks were aligned and processed using the TrakEM module in Fiji (Cardona et al., 2012).

Z-Stack of ultrastructural images (6nm pixel size in x-y and 100nm in z) was cropped and reduced in scale using Fiji and contrast was enhanced automatically. Segmentation of cells was done using Amira software (Thermo Fisher Scientific) (voxel size of 1-2-8.3). Cells within the cell cluster were outlined separately in x-y plane by using the segmentation tool and interpolated in z-plane. The segmented labels were visualized using the Volren display option.

## Acknowledgement

The authors thank the members of the Wittbrodt lab for critical feedback and support throughout the project. We are indebted to L. Centanin and G. Jékely for extensive and critical feedback on the manuscript. We thank the fish team (A. Sarazeno, M. Liv, E. Leist and R. Lipp) for excellent animal husbandry.

## Author contributions

C. S. and J. W. designed the experimental strategy, C. S. performed the experiments and in collaboration with X. P. and C. A. analyzed the data. R. C., I. W. and R. R. S. performed the EM analysis. C. S. created the display items with critical input from V. W. and L. Z.. C. S. and J. W. wrote the manuscript with refining input from all co-authors.

